# Deciphering inhibitory mechanism of coronavirus replication through host miRNAs-RNA-dependent RNA polymerase (RdRp) interactome

**DOI:** 10.1101/2022.06.18.496304

**Authors:** Olanrewaju B. Morenikeji, Muyiwa S. Adegbaju, Olayinka S. Okoh, Asegunloluwa E. Babalola, Anastasia Grytsay, Olubumi A. Braimah, Mabel O. Akinyemi, Bolaji N. Thomas

## Abstract

Despite what we know so far, Covid-19, caused by SARS-CoV-2 virus, remains a pandemic that still require urgent healthcare intervention. The frequent mutations of the SARS-CoV-2 virus has rendered disease control with vaccines and antiviral drugs quite difficult and challenging, with newer variants surfacing constantly. There is therefore the need for newer, effective and efficacious drugs against coronaviruses. Considering the role of RNA dependent, RNA polymerase (RdRp) as an important enzyme necessary for the virus life cycle and its conservation among coronaviruses, we investigated potential host miRNAs that can be employed as broad-range antiviral drugs averse to coronaviruses, with particular emphasis on BCoV, MERS-CoV, SARS-CoV and SARS-CoV-2. miRNAs are small molecules capable of binding mRNA and regulate expression at transcriptional or translational levels. Our hypothesis is that host miRNAs have the potential of blocking coronavirus replication through miRNA-RdRp mRNA interaction. To investigate this, we downloaded the open reading frame (ORF 1ab) nucleotide sequences and used them to interrogate miRNA databases for miRNAs that can bind them. We employed various bioinformatics tools to predict and identify the most effective host miRNAs. In all, we found 27 miRNAs that target RdRp mRNA of multiple coronaviruses, of which three - hsa-miR-1283, hsa-miR-579-3p, and hsa-miR-664b-3p target BCoV, SARS-CoV and SARS-CoV-2. Additionally, hsa-miR-374a-5p has three bovine miRNAs homologs viz bta-miR-374a, bta-miR-374b, and bta-miR-374c. Inhibiting the expression of RdRp enzyme via non-coding RNA is novel and of great therapeutic importance in the control of coronavirus replication, and could serve as a broad-spectrum antiviral, with hsa-miR-1283, hsa-miR-579-3p, and hsa-miR-664b-3p highly promising.

## Introduction

The diseases caused by SARS-CoV-2, a member of the Coronaviridae family, have had profound impact on all human endeavors, leaving hardship, death and destruction in its trail (Aftab et al 2020). The rate of transmission of SARS-CoV-2 from person to person is the major driver of the significant morbidity and mortality attendant to Covid-19 and its pandemic form (Gao et al., 2020; Walls et al., 2020). After a successive entry into the host, viral replication is another important step to its pathogenicity and transmission. A large portion of coronavirus genome encodes open reading frame (ORF) 1a/1b (Figure 1), which produces two precursor polyproteins (pp1a) and (pp1ab), dedicated to code for multiple enzymes among which is RNA dependent, RNA polymerase (RdRp). Each of this precursor polyproteins are subsequently cleaved into non-structural proteins (nsp). The pp 1ab is cleaved into 16 nsps, of which nsp12 or RNA dependent, RNA polymerase (RdRp) is one, and pivotal for successful virus replication in the host (Gao et al 2020). In addition, formation of protein complex between RdRp protein, nsp 7 and nsp 8 have been reported, as the latter duo serve as cofactor for RdRp (Kirchdoerfer and Ward 2019; Gao et al 2020). Except for retroviruses, most RNA viruses require the activity of RdRp protein for viral replication and may explain why its active site is the most conserved among these viruses (Aftab et al., 2020), thereby making it a prominent target for drug development.

**Figure 1.**
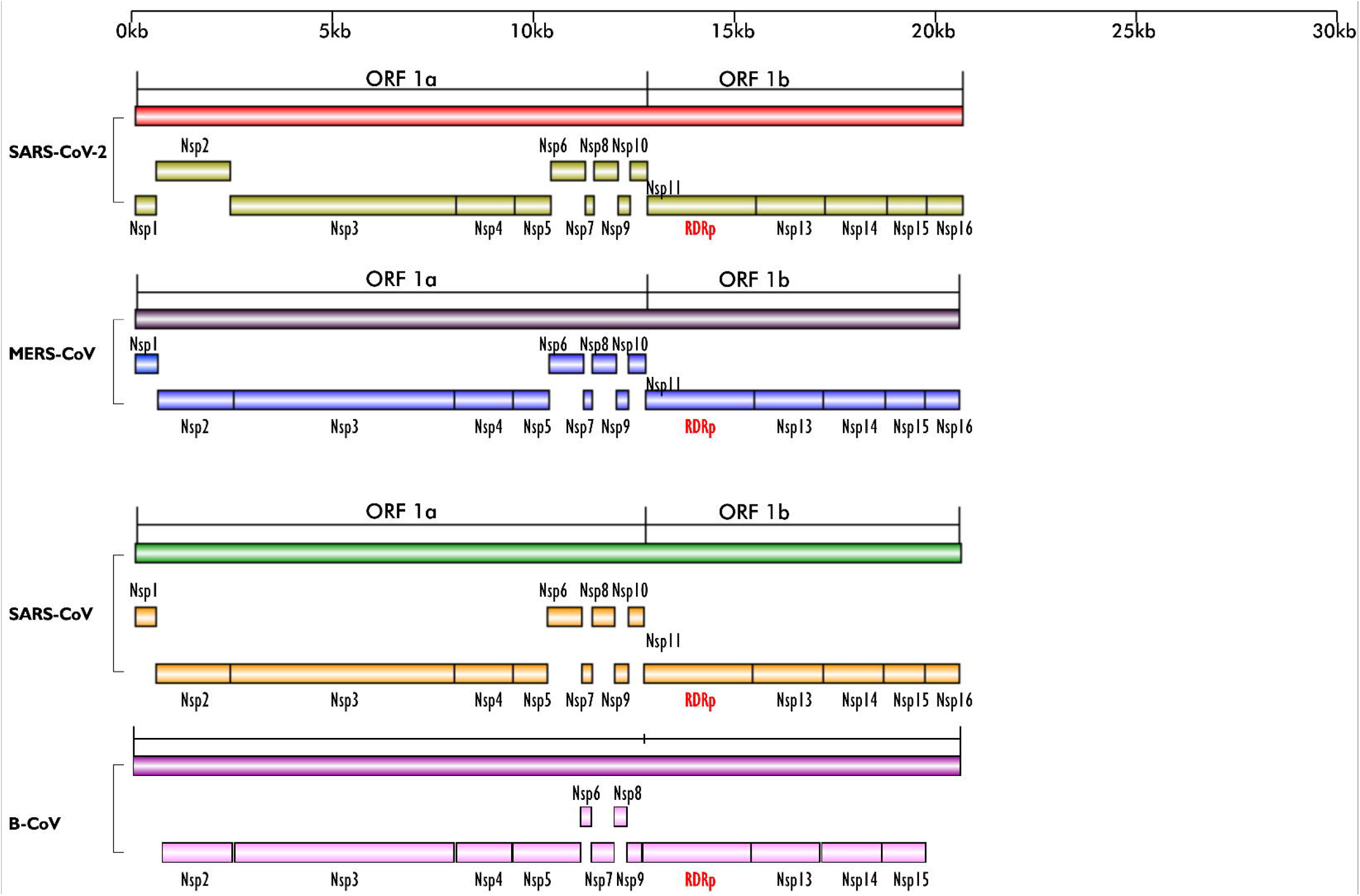
Schematics of the ORF 1a and ORF 1b regions of the genomes of SARS-CoV-2, MERS-CoV, SARS-CoV and BCoV; their encoded non-structural proteins (nsp) and RdRp layered on one another for easy comparison. The region is highly conserved in the four viruses; except for BCoV which does not have nsp1. All the nsps are present in the four viruses and they are arranged in the same sequence/order. Figures created with sketch pad,

Several vaccines have now been developed and approved for use to limit COVID-19 infection in humans. However, their safety and long-term efficacy against SARS-CoV-2 is not guaranteed (Saha *et al.*, 2021). Other strategies recommended for treating disease include inhibition of RdRp activity using antiviral agents like the nucleoside analogues, Favipiravir, Galidesivir, and Remdesivir, and other plant-based compounds such as Tellimagrandin I, Saikosaponin B2, Hesperidin and Epigallocatechin gallate (Saha et al., 2021). So far, these antiviral drugs have been reported to be ineffective against SARS-CoV-2, possibly due to single nucleotide polymorphism (SNP)-induced changes culminating in conformational, structural and functional amino acids changes and the high virus mutation rate. Therefore, alternate therapeutic options that are effective against the virus must be explored. Here, we propose an alternative option that utilizes blocking RdRp transcript via host microRNAs thereby inhibiting translation of the most important protein for viral replication, leading to reduced viral propagation and pathogenicity (Fig 2).

**Figure 2.**
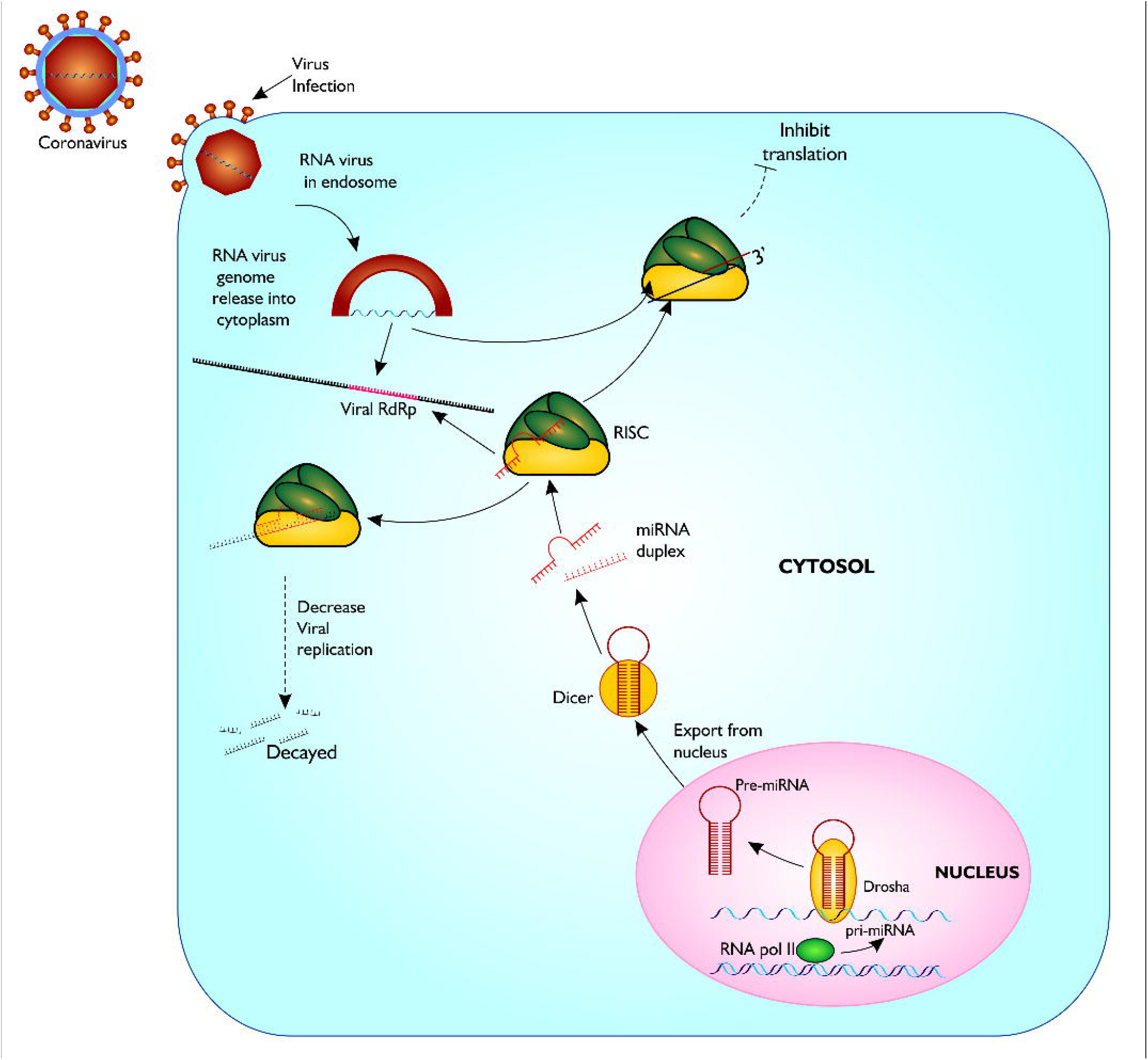
Proposed model of miRNA biogenesis and base pairing with coronavirus RdRp mRNA sequence. The figure gives a description of coronavirus infection of host, and release of host miRNA to base pair and degrade the virus or inhibit translation. Figure created with sketch pad.

MicroRNAs are short non-coding RNAs, of about 23 nucleotides in the introns (Trobaugh and Klimstra, 2017). They control several cellular operations transcriptionally by taking on target transcripts such as host mRNA and RNAs from the genome of pathogens, via sequence-specific interlink, influencing the function and/or stability of these targets (Morenikeji et al., 2020; Tucker et al., 2021). Several studies have shown the involvement of miRNAs in regulation of host immune responses. Morenikeji et al. (2020) demonstrated via *in silico* analysis that certain bovine miRNAs are involved in regulating specific immune response genes associated with Bovine coronavirus (BCoV) infection and were identified as drug targets and diagnostic biomarker for the virus. Additionally, miRNAs have been found in chicken binding the L gene region of Newcastle Disease virus, causing viral degradation and inhibiting replication *in vitro* (Chen et al., 2020). In humans, decrease in viral replication, translation and transmission from person to person due to binding of certain miRNAs to the genome of viruses such as Human immunodeficiency virus (HIV), Enterovirus 71 (E 71) and Human T cell leukemia virus, type I (HTLV-1) have also been reported (Nathans et al., 2009; Zheng et al 2013; Bai and Nicot 2015). More evidence on the involvement of miRNA in altering viral replication and pathogenesis have continued to emerge (Khongnomnan et al., 2015; Ingle et al., 2015; Trobaugh and Klimstra, 2017), but none of these studies have examined the role of miRNA in inhibiting coronavirus replication, showing the importance of our study.

Considering the significant role of RdRp in viral replication and survival, we elucidated host miRNAs that can bind mRNA of RdRp in four coronaviruses, resulting in its disintegration, thereby controlling the replication and pathogenesis of RNA viruses and opening a new door to therapeutic targets for coronaviruses.

## Materials and Methods

### Sequence mining of RdRp region from the genome of various coronaviruses

In this study, the analytical pipeline employed, starting from sequence curation to interactome networks, is a slight modification of our previously described model (Fig 3), (Morenikeji at al., 2020; 2021; Tucker et al., 2021). Since RdRp is one of the 16 non-structural proteins encoded by ORF 1ab gene of coronaviruses, we carried out an extensive search for ORF 1ab gene, and the nucleotide sequence of 13 selected coronaviruses, whose genomes have either been fully or partially annotated, were retrieved. These viruses are: SARS-CoV (NC_004718.3), SARS-CoV-2 (NC_045512.2), tylonycteris bat coronavirus (NC_009019.1), MERS-CoV (NC_019843.3), duck coronavirus (NC_048214.1), Canada goose coronavirus (NC_046965.1), BCoV (NC_003045.1), betacoronavirus England 1 (NC_038294.1), alphacoronavirus (NC_046964.1), bat coronavirus (NC_034440.1), pipistrellus bat (NC_009020.1), rabbit coronavirus (NC_017083.1), rodent and coronavirus (NC_046954.1)

**Figure 3.**
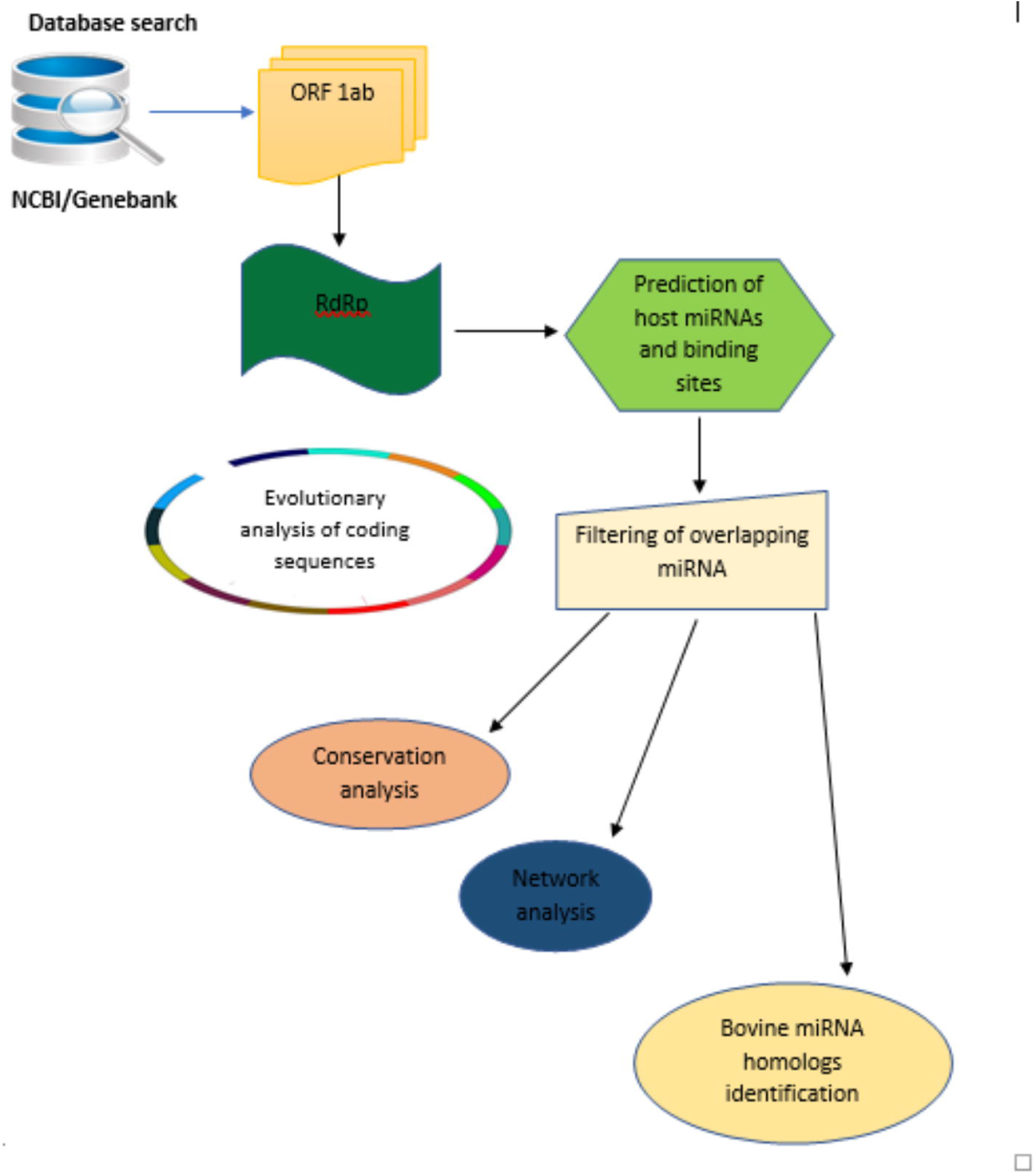
Flow chart of methodology used for the study. Step by Step pipeline for elucidating host miRNA – viral RdRp interaction.

### Evolutionary analysis of RdRp in 13 coronaviruses

To determine the evolutionary relationship and distance of the RdRp region from the 13 coronaviruses, we constructed a phylogenetic tree using https://ngphylogeny.fr/ with the following workflow. Preliminary multiple sequence alignment (MSA) was generated using MAFFT, followed by trimming of the MSA to focus on the informative regions using block mapping and gathering with entropy (BMGE) (Criscuolo et al., 2010). The phylogenetic tree was inferred using PhyML (Guindon et al., 2010) and tree visualization carried out with interactive tree of life (iTol) (https://itol.embl.de). Using the MSA generated by BMGE, pairwise distance between the RdRp of the 13 coronaviruses was computed using MEGA X (Kumar et al., 2018).

### Prediction and network of miRNA binding sites in the RdRp region of MERS-CoV, BCoV, SARS-CoV and SARS-CoV-2

To examine whether host cellular miRNA can target coronavirus RdRp, we selected four common coronaviruses in human and cattle for further analysis. Potential miRNA binding sites in the RdRp coding regions of MERS-CoV, BCoV, SAR-CoV and SARS-Cov-2 were predicted using mirDB software (http://mirdb.org). Each of the RdRp coding sequence from the 4 coronaviruses were used as the target sequence and human genome selected as the reference genome for miRNA prediction. After each prediction, miRNAs with a score of 60 and above were considered significant and selected for further analysis. The list of miRNAs from each coronavirus were intersected with Bioinformatic and Evolutionary Genomics (BEG) Venn diagram generator (http://bioinformatics.psb.ugent.be/webtools/Venn/). Based on the complementary base pairing of miRNAs and RdRp mRNA and the value of normalized binding free energy (ndG), possible miRNA-mRNA interactome network connections were determined using Cytoscape version 3.7.2, as previously described (Morenikeji et al., 2020; Tucker et al., 2021). To search for possible homologs of human miRNAs in the bovine genome, we searched the miRNA database (https://mirbase.org). The sequence of each of the top 25 miRNAs selected were used as query against the *Bos taurus* genome on used mirDB. Homologous bovine miRNAs were extracted and recorded for further analysis.

## Results

### Dataset of RdRp nucleotides from the genome of 13 coronaviruses

Our search for nucleotide sequences encoding for RdRp in coronaviruses using the keyword “1ab polyprotein” initially yielded about 59 organisms. After filtering for only coronaviruses, 13 viruses, whose genome was either fully or partially annotated were subsequently selected for further analysis. The RdRp encoding regions of tylonycteris bat coronavirus, MERS-CoV, duck coronavirus, SARS-CoV, Canada goose coronavirus, BCoV, betacoronavirus England 1, alphacoronavirus, bat coronavirus, SARS-CoV-2, pipistrellus bat, rabbit coronavirus and rodent coronavirus were identified to be within the ORF 1ab gene (Table 1). Since RdRp protein is categorized to be one of the cleaved 16 non-structural proteins encoded by ORF 1ab gene, its coding region which falls between nsp11 and nsp 13, is beyond the coding region of ORF 1a gene, which partially overlaps with ORF 1ab gene and encode variants of nsp1 to nsp9 (Fig 1). Each of the RdRp nucleotides for the 13 coronaviruses were copied and added to the pipeline (Fig 3) to determine their evolutionary relationship.

**Table 1.** List of coronaviruses; accession number of their ORF1ab gene, genome location and the location of RdRp coding sequence within ORF 1ab genome location.

| S/N | Virus | Accession Number | Genome Location of ORF 1ab | Location of RdRp within ORF 1ab |
| --- | --- | --- | --- | --- |
| 1 | Tylonycteris bat CoV | NC_009019.1 | 267-21625 | 13553-16327 |
| 2 | SARS-CoV | NC_004718.3 | 265..21485 | 13401-16163 |
| 3 | MERS-CoV | NC_019843.3 | 279-21514 | 13410-16207 |
| 4 | Duck coronavirus | NC_048214.1 | 347..20364 | 12211-15071 |
| 5 | Bovine coronavirus | NC_003045.1 | 211..21494 | 13318-16100 |
| 6 | Canada goose CoV | NC_046965.1 | 554..20085 | 11971-14786 |
| 7 | Betacoronavirus England 1 | NC_038294.1 | 278-21513 | 13400-16185 |
| 8 | Alphacoronavirus Bat-CoV | NC_046964.1 | 281-20175 | 12136-14885 |
| 9 | Bat CoV | NC_034440.1 |  | 13156-15970 |
| 10 | Pipistrellus bat coronavirus | NC_009020.1 | 261-21808 | 13661-16332 |
| 11 | Rabbit coronavirus | NC_017083.1 | 209-21663 | 13483-16270 |
| 12 | Rodent coronavirus | NC_046954.1 | 211-21596 | 13386-16201 |
| 13 | SARS-CoV-2 | NC_045512.2 | 266-21555 | 13430-16221 |

### Evolutionary relatedness of RdRp coding sequences

To confirm that RdRp coding sequence is conserved among the coronaviruses, we examined their evolutionary relatedness. The sequence analysis of RdRp reveal minor variation across the 13 viruses though sharing common evolutionary origin (Fig 4). Two viruses, MERS-CoV and Betacoronavirus England 1, are not different from each other in this region, showing a pairwise distance of 0.00 (Table 2). Similarly, comparing the RdRp sequences of bat coronavirus with either MERS-CoV or Betacoronavirus England 1 indicated some level of closeness with a value of 0.18. A similar close relatedness was observed for SARS-CoV and SARS-CoV-2 having a pairwise distance of 0.29. Alphacoronavirus is the most distantly related from the rest of the viruses, showing consistent higher value for the pairwise distance, further supported by the phylogenetic tree analysis(Fig 4).

**Figure 4.**
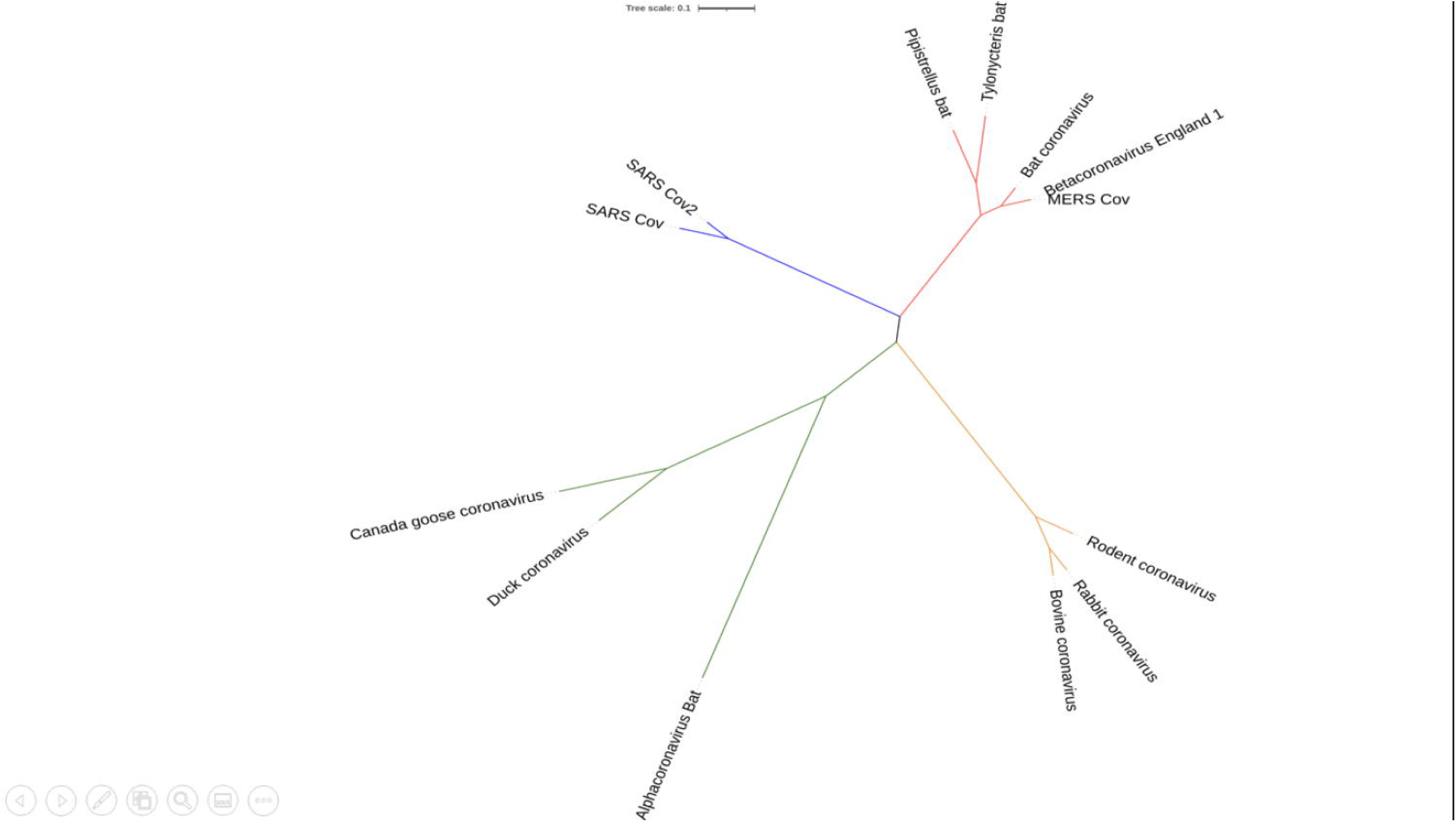
Phylogenetic tree showing the evolutionary relationship of 13 coronaviruses. Tree was constructed using MEGA X.

**Table 2.**
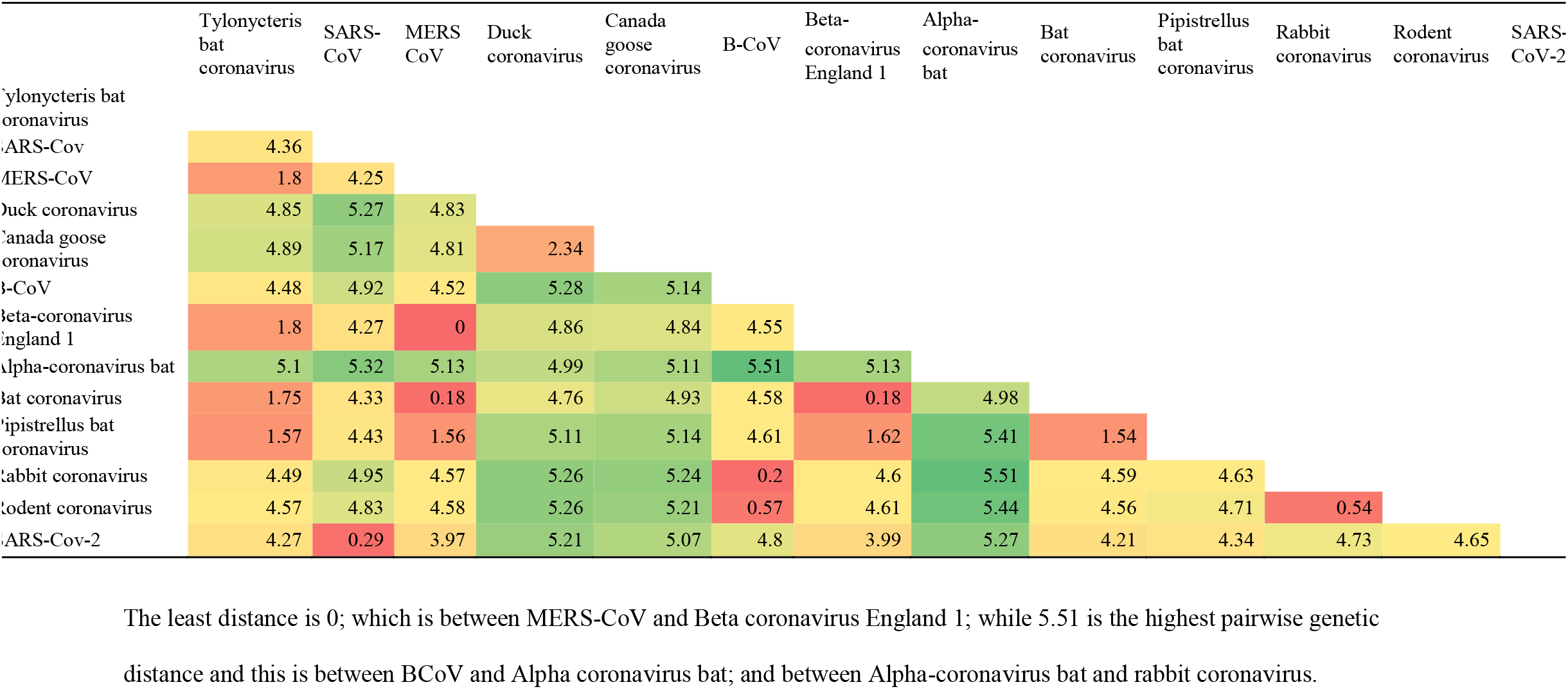
Genetic pairwise distance of the 13 coronaviruses used in the study. The least distance is 0; which is between MERS-CoV and Beta coronavirus England 1; while 5.51 is the highest pairwise genetic distance and this is between BCoV and Alpha coronavirus bat; and between Alpha-coronavirus bat and rabbit coronavirus.

### ORF 1 ab is conserved in BCoV, MERS-CoV, SARS-CoV and SARS-CoV-2

From the genome organization of BCoV, MERS-CoV, SARS-CoV and SARS-CoV-2 (Fig 1), ORF 1ab in the viruses are very similar, indicating high level of conservation in this gene making it an excellent antiviral drug candidate. To ascertain the level of conservation of ORF 1ab in BCoV, MERS-CoV, SARS-CoV and SARS-CoV-2, the well annotated ORF 1ab genes of each of the viruses were overlayed on one another and compared. Comparing the gene across the viruses, we found that ORF 1ab is highly conserved across the viruses as no conspicuous difference was noted in the arrangement of all the non-structural proteins and RdRp (Fig 1), making RdRp a good antiviral drug target.

### Identification of miRNA binding to RdRp of BCoV, MERS-CoV, SARS-CoV and SARS-CoV-2

In this analysis, BCoV, MERS-CoV, SARS-CoV and SARS-CoV-2 miRNAs that bind to the RNA-dependent RNA polymerase (RdRp) of coronaviruses were examined. A total of one hundred and three (103) miRNA were obtained for BCoV, seventy-eight (78) for MERS-CoV, fifty-seven (57) for SARS-CoV, while ninety-seven (97) miRNAs were found for SARS-CoV-2. To ensure the binding of miRNAs to RdRp target, significant miRNAs were filtered based on the ranking scores as described above. The filtering generated a total of sixty-six (66) miRNA for BCoV, forty-one (41) for MERS-CoV, twenty-nine (29) for SARS-CoV and fifty-three (53) for SARS-CoV-2 (Fig 5). The results of complementary binding of human miRNAs to the RdRp sequence for each of the 4 coronaviruses were intersected to identify broad-spectrum miRNAs, which can possibly inhibit viral replication. As shown, there was no miRNA that could bind to the RdRp region of all 4 viruses (Fig 6a). However, we uncovered three miRNAs; hsa-miR-1283, hsa-miR-664b-3p and hsa-miR-579-3p that could bind to this region in BCoV, SARS-CoV and SARS-CoV-2 (Table 3; Fig 6b). Similarly, miRNAs that could bind to the region in at least two coronaviruses were identified, ranging from as low as one (hsa-miR-8081) for MERS-CoV and SARS-CoV to as high as nine (hsa-miR-585-5p, hsa-miR-7159-5p, hsa-miR-1305, hsa-miR-15a-5p, hsa-miR-6507-5p, hsa-miR-16-5p, hsa-miR-3065-5p, hsa-miR-195-5p and hsa-miR-15b-5p) for BCoV and MERS-CoV (Table 3).

**Figure 5.**
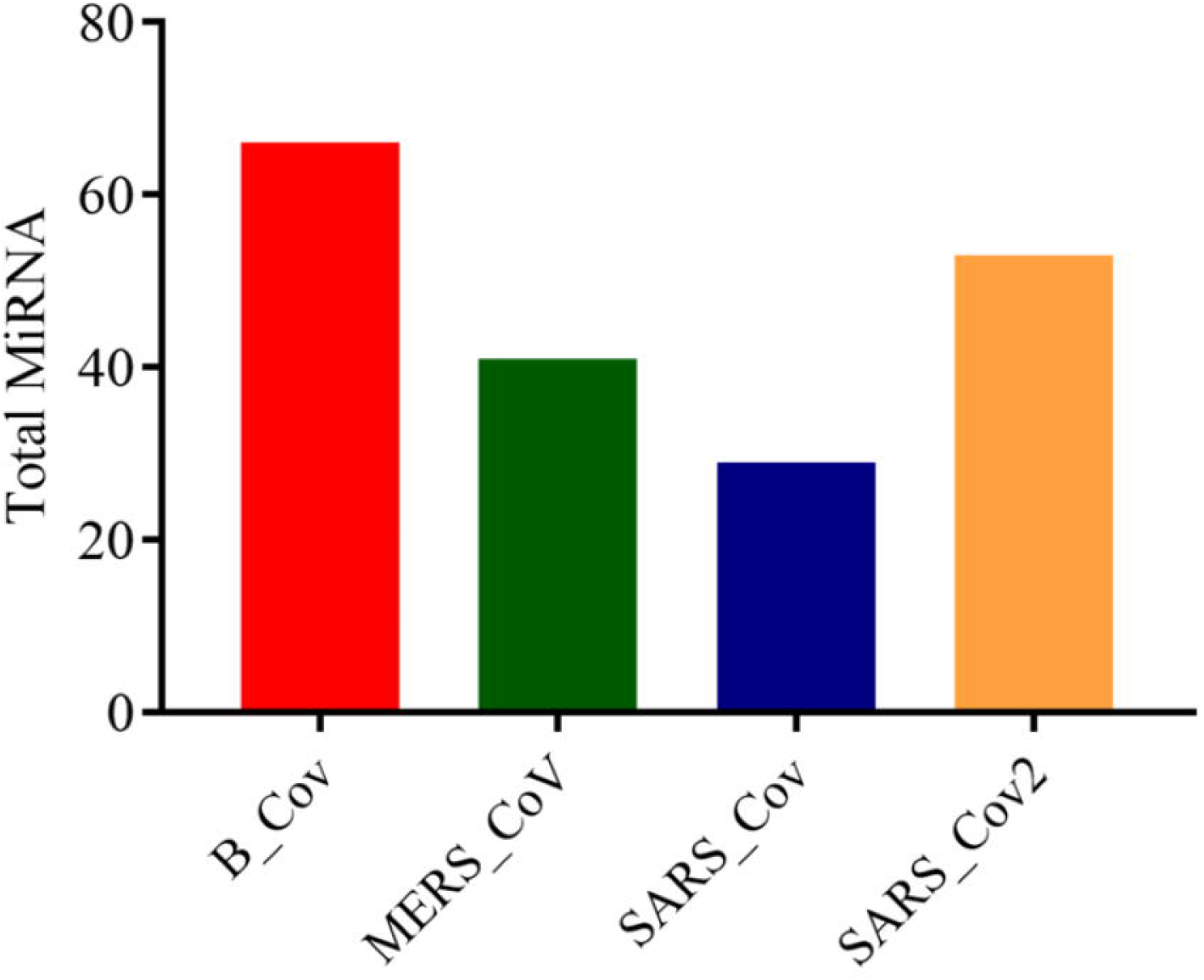
Bar chart showing the number of predicted human miRNA that can bind with the ORF 1ab region of each of BCoV, MERS-CoV, SARS-CoV and SARS-CoV-2. Figure created graphpad

**Figure 6.**
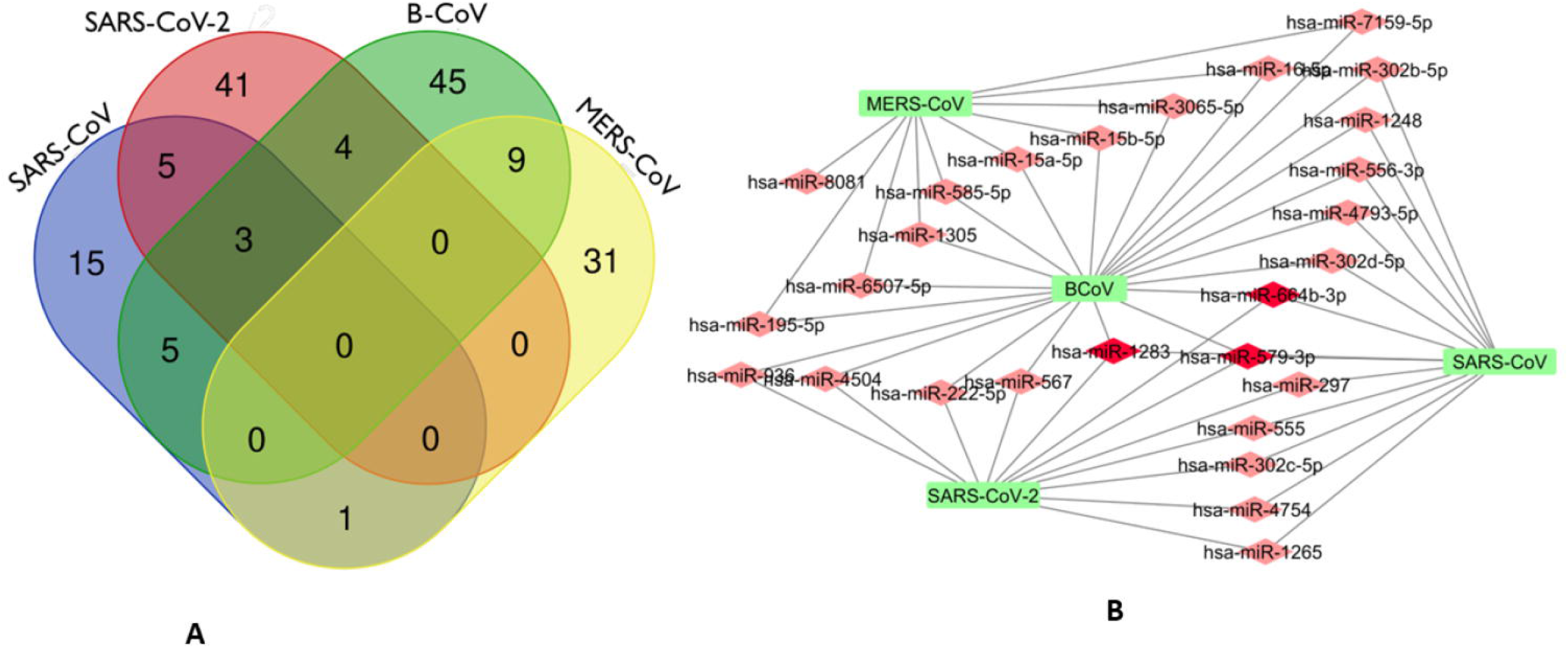
Venn Diagram (http://bioinformatics.psb.ugent.be/webtools/Venn/) showing the number of predicted human miRNA that can target multiple coronaviruses. The number in the intersection/overlapping regions represent the number of miRNAs that can concomitantly target the coronaviruses represented by the intersected shape. (a). Network connections among miRNAs and RdRp of SARS_CoV, B_CoV, MERS_CoV and SARS_CoV-2. B. Generated using Cytoscape 3.7.2

**Table 3.**
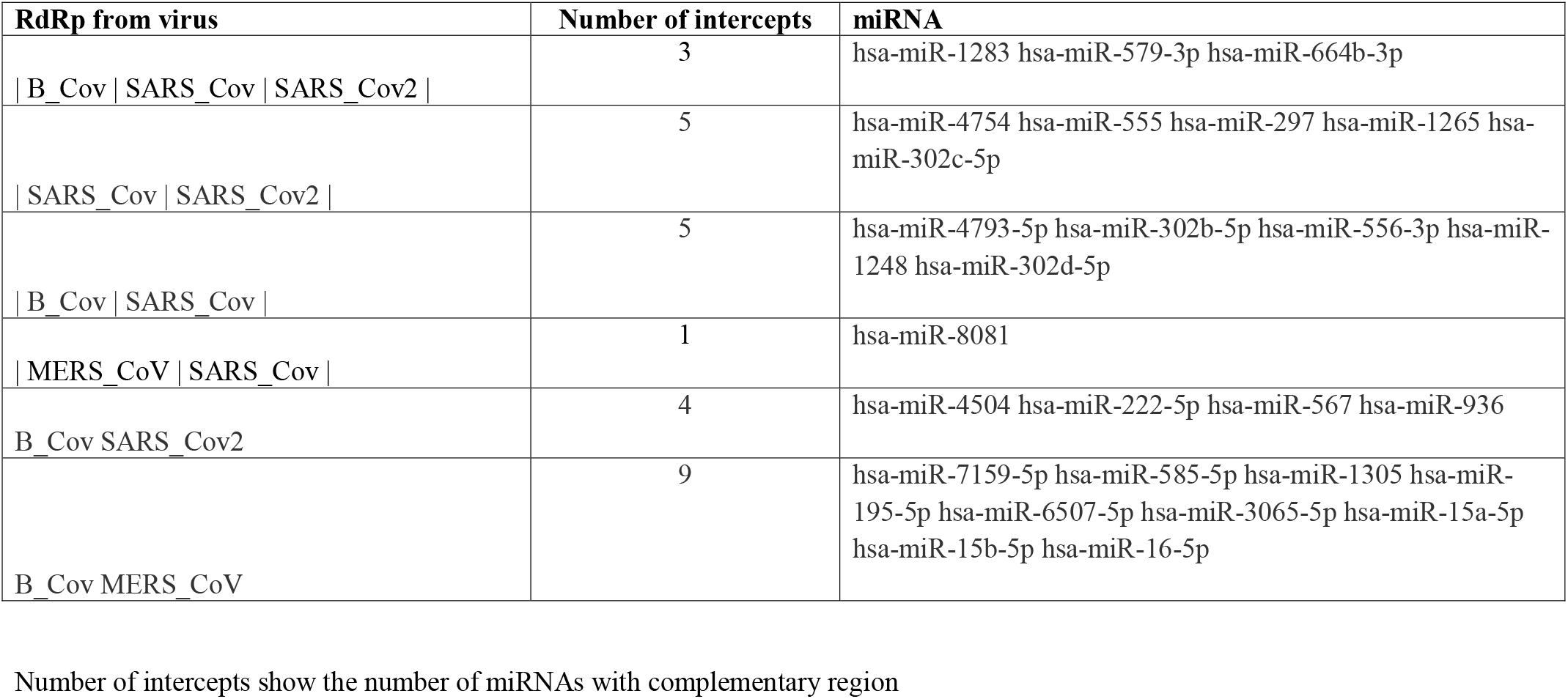
Predicted miRNAs with regions of complementarity in multiple coronaviruses from BCoV, MERS-CoV, SARS-CoV and SARS-CoV-2. Number of intercepts shows the number of miRNA that have complementary region, while the miRNAs are listed under the MiRNA column.

Interestingly, there was no miRNA concomitantly binding RdRpregion in both MERS-CoV and SARS-CoV-2 (Fig 6a). The identity of the connections of miRNAs and RdRp between for coronavirues (BCoV, MERS-CoV, SARS-CoV and SARS-CoV-2) are depicted through a network as shown in Figure 6b. This interactome reveal a possible molecular mechanism for regulating multiple coronavirus replication through miRNAs binding RdRp mRNA. Network of different nodes were created based on all identified miRNAs and potential binding sites on RdRp mRNA, while the network edges were determined through the value ndGs and correlation between each RNA. From Table 3, it is shown that 27 human miRNAs targeted multiple viruses from BCoV, MERS-CoV, SARS-CoV and SARS-CoV-2. Of particular importance, among these 27 miRNAs are three miRNAs (hsa-miR-1283, hsa-miR-579-3p and hsa-miR-664b-3p) that are predicted to target BCoV, SARS-CoV and SARS-CoV-2. Furthermore, we report five miRNAs targeting SARS-CoV and SARS-CoV-2, while another five targeted BCoV and SARS-CoV.

### Human miRNAs homologs found in bovine genome target RdRp mRNA sequences

Of the top 25 human miRNAs selected for further analysis, eight has bovine miRNA homologs as shown (Table 4). Only one of the human miRNAs, hsa-miR-374a-5p, had three bovine miRNA homologs including bta-miR-374a, bta-miR-374b, and bta-miR-374c. hsa-miR-3065-5p has two bovine homologs - bta-miR-338 and bta-miR-3065, while others have one homolog each. In all, we report 13 bovine homologs, and two of them, bta-miR-196a and bta-miR-338 are read in reverse direction while eleven are forward stranded.

**Table 4.** Predicted human miRNAs that have bovine miRNA homologs, their size and strands. The seed location is with respect to the human miRNA, while the size and strand are the bovine miRNAs. – strand miRNAs are read in reverse direction

| miRNA | Seed location | Bovine homolog | Size | Strand |
| --- | --- | --- | --- | --- |
| hsa-miR-196a-1-3p | 229, 1027, 1413, 1765, 1940 | bta-miR-196a | 3 to 20 | - |
| hsa-miR-654-5p | 1387 | bta-miR-380-5p | 1 to 22 | + |
| hsa-miR-541-3p | 1387 | bta-miR-541 | 1 to 22 | + |
| hsa-miR-374a-5p | 1047, 1362, 1370 | bta-miR-374a | 1 to 22 | + |
|  |  | bta-miR-374b | 2 to 22 | + |
|  |  | bta-miR-373c | 1 to 21 | + |
| hsa-miR-664b-3p | 1628, 2626 | bta-miR-664b | 3 to 23 | + |
| hsa-miR-545-5p | 830, 2750 | bta-miR-545-5p | 1 to 21 | + |
| hsa-miR-374b-5p | 1047, 1362, 1370 | bta-miR-374b | 1 to 22 | + |
|  |  | bta-miR-374a | 1 to 22 | + |
|  |  | bta-miR-374c | 1 to 21 | + |
| hsa-miR-3065-5p | 227, 512, 1025 | bta-miR-338 | 1 to 23 | - |
|  |  | bta-miR-3065 | 1 to 23 | + |

## Discussion

The genome arrangement of coronaviruses is similar and of particular importance is the open reading frame 1ab (ORF 1ab) gene, which encodes 1ab polyprotein, a protein precursor that is further cleaved into sixteen non-structural proteins (nsps). One of the sixteen nsps is RdRp that plays a vital role in RNA virus replication (Aftab et al., 2020; Gao et al 2020; Jiang et al., 2021). RdRp is a promising candidate for drug target for treating diseases caused by coronavirus because the active site is highly conserved and the protein lacks homologous counterparts in host cell (Jiang et al., 2021). Medical interventions in form of mRNA vaccines and antiviral drugs have been developed and approved to treat coronavirus disease such as Covid-19, but many of these drugs are still undergoing clinical trials. The concept behind antiviral drugs for treating Covid-19 and other diseases caused by other RNA viruses is to identify compounds which can bind to active site of the RdRp enzymes and prevent its catalytic activity, which leads to viral replication (Yang et al., 2011; Markland et al., 2000; Elfiky 2016; Elfiky et al., 2017; Elfiky and Ismail 2019; Ezat et al., 2019; Wang et al., 2020). However, major concern on the efficacy of the antiviral drugs remains, necessitating exploring alternative options, including preventing RdRp protein translation. To the best of our knowledge, the use of non-coding RNA such as miRNAs as an alternative route has not been explored as antiviral drug option. Since miRNA can bind directly to the genome of RNA virus or cause changes in the host transcriptome facilitated by the virus, it is noteworthy that finding host miRNA that can bind directly to the RdRp region of coronaviruses may provide insight on effective manipulation to control the viral replication/load in the host and provide a remarkable alternative treatment.

To have an effective antiviral drug, it is important that a virus target be conserved. Therefore, we identify the conservation of miRNA binding site in the RdRp sequence of multiple coronaviruses through evolutionary analysis. First, 13 annotated RdRp nucleotide sequences were used to define the conservation of this region and construct a phylogenetic tree. Most of the viral species examined belong to the betacoronavirus subfamily, with our result showing high similarity between them, given this region as a potential drug target. Interestingly, the pairwise distance result between MERS-CoV and Betacoronavirus England 1 show no difference in this region for both viruses, indicating that they are likely to have the same binding site for host miRNAs. A close similarity in the RdRp region of SARS-CoV and SARS-CoV-2 shows the virus evolved from a common origin, in agreement with previous findings (Wu et al 2020; Aftab et al 2020). The genetic conservation of RdRp gene across multiple viruses shows a strong positive selection for this region and justifies the fact that the enzyme coded by this region is important for almost all RNA virus replication, strengthening its choice for miRNA drug targeting. It is puzzling that the magnitude of the difference in pathogenicity, rate of transmission and virulence between SARS-CoV and SARS-CoV-2 is only caused by single nucleotide mutations (Ceraolo and Giorgi, 2020; Lu et al., 2020; Kruse, 2020; Nguyen et al., 2020; Wang et al., 2020). Thus, the slight pairwise distance observed between the RdRp sequences of SARS-CoV and SARS-CoV-2 may have remarkable implications on the number of miRNA which can bind concomitantly with both viruses. Several regions of RNA viruses mutate at a faster rate as a mechanism to escape host immune system reaction, however, a slower mutation rate at the RdRp region means a miRNA could be broad spectrum antiviral drug for many viruses.

Remarkably, our study uncovered several miRNAs that bound to the RdRp sequence of coronaviruses including SARS-CoV, SARS-CoV-2 MERS-CoV and BCoV. The presence of human miRNA homologs in bovine genome is of great importance as this is indicative of the crucial roles these miRNAs play as preserved by evolutionary forces or selection. Additionally, some of the miRNAs have multiple binding sites within the RdRp region thereby increasing the binding probability and reducing off-target effects, strengthening their choice for possible antiviral molecule. This is contrary to the submission of Thorg et al. (2017), which stated that the common location of the miRNA binding site is the UTRs of the viral genome. Conversely, our results align with similar study in chicken where Wang et (2021) identified multiple miRNA binding sites within the L gene of the NDV and infectious bursal disease virus (IBDV). Wang et al (2021) found the overexpression of ggam-miR-21 inhibiting VP1 translation in chicken fibroblasts and suppresses overall viral replication.

We identified some miRNAs including hsa-miR-1283, hsa-miR-664b-3p, hsa-miR-579-3p, which targeted multiple regions in the RdRp sequence of BCoV, SARS-CoV, and SARS-CoV-2, these miRNAs have been previously linked with onco-protective roles, indicating their growth regulatory function. For example, hsa-miR-1283 has been linked with cardio-protection and inhibition of apoptosis (Liu *et al.*, 2021), thereby blocking oncogenesis. In addition, this miRNA has also been implicated in hypertension (Chen *et al.*, 2021). hsa-miR-579-3p, on the other hand has been reported to be associated with growth control and tumor suppression via control of melanoma progression (Fattore *et al.*, 2016; Kalhori et al., 2019), while hsa-miR-664b-3p is reported to play a critical role in regulating cancer progression (Liu *et al.*, 2020). From our study, an upregulation or administration of any of the three miRNAs might play a dual role of blocking viral replication/degradation and inhibition of cancer progression.

## Conclusion

In summary, we utilized several computational approaches to examine genome plasticity and elucidate potential host miRNAs that could bind to the RdRp sequence region of coronaviruses. Although viral genome is known to be variable, we report high conservation of RdRp sequence in multiple coronaviruses species, indicating evolutionary favorability, hence a candidate signature for genome targeting. In all, this study also provides an insight on possible alternative route for targeting and inhibiting viral replication via host non-coding RNA (miRNAs) to combat disease rather than common anti-coronavirus drug, based on inhibiting RdRp enzymatic activities. In particular, hsa-miR-1283, hsa-miR-664b-3p, hsa-miR-579-3p and hsa-miR-374a-5p with bovine homologs bta-miR-374a, bta-miR-374b, and bta-miR-374c are very promising. This study opens the door for developing non-coding RNAs as a broad-spectrum antiviral therapy and lays a foundation for further investigation to validate the effective binding of identified miRNAs to RdRp sequences of coronaviruses through *in vivo* or *in vitro* analysis.

## Acknowledgement

OBM was supported by Pitt-Momentum Fund of the University of Pittsburgh. Ongoing support by the Division of Biological and Health Sciences, Pitt-Bradford, College of Health Sciences and Technology, Rochester Institute of Technology is acknowledged APC charges for this article were fully paid by the University Library System, University of Pittsburgh.

## Author Contributions

OBM conceptualized and designed the experiments; MSA, OSO, AEB and OBM carried out the experiments, analyzed the data and drafted the manuscript; MSA, OSO, AG, OAB, MOA, BNT and OBM revised the manuscript, contributed to the discussion and scientific content. All authors read and approved the final version of the manuscript.

## Data Availability (see supplementary Tables)

